# Inhibition of Osteoclast Differentiation and Activation by Leukotriene B4 Loaded in Microspheres

**DOI:** 10.1101/2020.05.22.111633

**Authors:** Francine Lorencetti-Silva, Maya Fernanda Manfrin Arnez, João Pedro de Queiroz Thomé, Fabrício Kitazono de Carvalho, Lúcia Helena Faccioli, Francisco Wanderley Garcia Paula-Silva

## Abstract

**Aim:** Leukotriene B_4_ (LTB_4_) is a labile inflammatory lipid mediator important for host defense. We hypothesised that sustained delivery of LTB_4_ would be a therapeutic strategy to prevent osteoclast cell differentiation in bone resorption in inflammatory diseases. Therefore, the aim of this study was to investigate the role of LTB_4_ in differentiation of monocytic lineage cells into osteoclasts after stimulation with LTB_4_ loaded in microspheres (MS).

**Design:** LTB_4_-MS were prepared using an oil-in-water emulsion solvent extraction-evaporation process. Sterility, LPS contamination, characterization and efficiency of LTB_4_ encapsulation were investigated. J774A.1 cells were cultured in the presence of monocyte colony stimulating factor (M-CSF) and ligand for receptor activator of nuclear factor kappa B (RANKL) and then stimulated with LTB_4_-MS. Cytotoxicity was determined by lactate dehydrogenase assay, osteoclast formation by means of the activity of tartrate-resistant acid phosphatase enzyme and gene expression was measured by quantitative reverse transcription polymerase chain reaction to investigate regulation of *Alox5, Alox5ap, Acp5, Mmp9, Calcr* and *Ctsk*.

**Results:** We found that 5-lipoxygenase pathway is involved in the osteoclastic differentiation hematopoietic lineage cells and that exogenous addition of LTB_4_-MS inhibited osteoclastogenesis induced by M-CSF and RANKL. The mechanism of LTB_4_-MS involved induction of *Mmp9* gene expression and inhibition of *Calcr* and *Ctsk*, without changing *Acp5*.

**Conclusion:** LTB_4_-MS inhibited differentiation of macrophages into an osteoclastic phenotype and cell activation under M-CSF and RANKL stimulus shedding light on a potential therapeutic strategy to prevent osteoclast differentiation.

## Introduction

Bone resorption resulting from periapical inflammation is a process that involves a series of steps, starting with the recruitment of monocytes from hematopoietic lineage, followed by osteoclast differentiation and maturation, ultimately resulting in bone loss (Tay et al., 2004; Gallagher, 2008; Paula-Silva et al., 2010; Trouvin and Goeb, 2010; Boyce, 2013).

Several cytokines and growth factors are essential for differentiation and maturation of osteoclasts, including interleukins (IL-1α, IL-6, IL-11, IL-15, IL-17 and IL-18), tumor necrosis factor-alpha (TNF-α), monocyte colony-stimulating factors (M-CSF, CSF2 and CSF3), prostaglandins and leukotrienes (Garlet et al., 2006; Fracon et al., 2008; Blackwell et al., 2010; Lee et al., 2012; McCoy et al., 2013; Lin and O’Connor, 2014). These factors stimulate osteoclast progenitor cells or in the regulation of a paracrine system that includes the molecules RANK (Receptor activator of nuclear factor kappa B), ligand for RANK (RANKL) and osteoprotegerin (OPG) (Hofbauer and Heufelder, 2001; Zhao et al., 2007; Boyle et al., 2003; Boyce, 2013).

The canonical process of osteoclastogenesis is regulated by soluble mediators RANKL and OPG. Briefly, RANK is a receptor expressed on the surface of osteoclasts and their precursors, dendritic cells and activated T cells (Boyle et al., 2003; Wang and McCauley, 2011; Boyce, 2013). RANKL is a soluble mediator, member of the tumor necrosis factor superfamily, synthesized by osteoblasts, bone marrow cells and endothelial cells, which induces osteoclast activation and fusion when it binds to the RANK (Sabeti et al., 2005; Leibbrandt and Penninger, 2008). RANKL induce the expression of osteclastogenic lineage genes including tartrate-resistant acid phosphatase (*Acp5*), cathepsin K (*Ctsk*), matrix-9 metalloproteinase (*Mmp9*) and the calcitonin receptor (*Calcr*), thus leading to maturation of these cells (Boyle et al., 2003; Lau et al., 2005; Boyce, 2013). Effects of RANKL are blocked by soluble receptors, such as osteoprotegerin, secreted by osteoblasts, which inhibits osteoclast differentiation (Jin et al., 2007; Raggatt and Partridge, 2007; Belibasakis et al., 2013; Kang et al., 2014). Overexpression of OPG blocks osteoclast maturation, leading to the development of osteopetrosis in rats, and deletion of the OPG gene results in increased bone resorption, inducing the development of osteoporosis (Kearns et al., 2008).

Previous studies have demonstrated that bacterial lipopolysaccharide (LPS) or the contamination of dental root canals induced periapical inflammation and the expression of the *Alox5* gene, responsible for encoding the 5-LO enzyme, concomitantly with the expression of *Tnfrsf11a, Tnfsf11* and *Tnfrsf11b* genes, which encode osteoclastogenesis mediators RANK, RANKL and OPG, respectively (Paula-Silva et al., 2016; Ribeiro-Santos et al., 2019; Paula-Silva et al., 2020). Signaling between cells of the immune system, cells of the hematopoietic lineage and resident cells might result in degradation of organic and inorganic matrices of bone tissue, depending on the nature of the mediators involved in the process (Tay et al., 2004; Gallagher, 2008; Paula-Silva et al., 2010; Trouvin and Goeb, 2010, Boyce, 2013).

Lipid mediators derived from arachidonic acid are widely studied in inflammation and inflammatory cell differentiation (Secatto et al., 2014; Zoccal et al., 2016; Prado et al., 2017; Pereira et al., 2018). However, the role of these mediators in osteoclast precursor cell populations present at the inflammation site is poorly understood. Therefore, the aim of this study was to investigate the role of leukotriene B4 in the differentiation of cells from the monocytic lineage (J774A.1) into osteoclasts, after stimulation with leukotriene B4 encapsulated in microspheres. This therapeutic strategy, based on the use of polylactic-co-glycolic acid microspheres (PLGA), promote a sustained delivery of lipid mediator over time since lipid mediators are extremely labile (Nicolete et al., 2009; Santos et al., 2011; Pereira et al., 2015; Reis et al., 2017; Lorencetti-Silva et al., 2019).

## Methods

### Preparation of microspheres

Microspheres (MS) were prepared as a pharmacological strategy using an oil-in-water emulsion solvent extraction-evaporation process (Nicolete *et al*., 2007; Nicolete *et al*., 2008). Briefly, LTB_4_ (CAYM-14010; Cayman Chemical Company, Michigan, USA) was dissolved in absolute ethanol (100 µg/mL). Then, 0.3 mL of the organic phase, equivalent to 3× 10^−5^M of the LTB_4_ solution was added to 10 mL of methylene chloride supplemented with 30 mg of 50:50 poly (lactic-co-glycolic acid) (PLGA) (Boehringer Ingelheim, Germany). Next, 40 mL of 3% polyvinyl alcohol (3% w/v PVA) (Sigma-Aldrich CO., St. Louis, MO, USA) were added and the mixture was mechanically stirred at 600 rpm for 4 h (RW-20; Ika®-Werke GmbH & CO. KG, Staufen, Germany). Microspheres were washed (3x) with deionized water (Milli-Q®, Merck Millipore, Darmstadt, Germany), lyophilized, and stored at −20 °C until use.

### Sterility and LPS contamination tests

A sterility test was performed in which small microsphere aliquots were diluted in 500 µL of 1x PBS (phosphate buffered saline) and 100 µL of solution was spread on Brain Heart Infusion (BHI)-Agar medium and kept in an incubator at 37° C for 24 h to detect microbial contamination.

Microspheres were tested for LPS contamination using the Limulus Amebocyte Lysate (LAL) QCL-1000™ kit (Lonza Walkersville, Inc., Olten, Switzerland) according to the manufacturer’s instructions. To obtain the standard curve, the serial dilution regime was adopted, starting from 1.0 EU / mL of *E. coli* endotoxin 0111: B4 (E50-640). Optical density was analyzed at a wavelength of 405 ηm in a μQuantTM spectrophotometer (BioTek® Instruments Inc., Winooski, USA), using the KC4™ Data Analysis Software (BioTek® Instruments Inc.), to determine the concentration of endotoxin units in each ml of solution containing microspheres (EU / ml).

### Characterization of microspheres

Size distribution of MS was determined using a LS 13 320 Laser Diffraction Particle Size Analyzer (Beckman Coulter, USA). Samples (1 mg) of either unloaded-MS or LTB_4_ -loaded MS was dispersed in 0.4 mL of purified sterile water and then analyzed at 25 °C. Zeta potential of MS was determined using a Zetasizer Nano (Malvern Instruments, England). Each sample was prepared dispersing 1 mg of unloaded-MS or LTB_4_-loaded MS in 0.4 mL of purified water containing 10 mM NaCl and then analyzed at 25 °C. Morphology of MS samples was assessed by scanning electron microscopy (SEM) using a FEI Inspect S 50 scanning microscope (FEI; Oregon, USA).

### Efficiency of LTB_4_ encapsulation in MS

For calculation of encapsulation efficiency, samples of LTB_4_-loaded MS (4 mg) were dissolved in 1 mL of acetonitrile/ethanol (7:3 v/v), to disrupt the MS structure. The solvent was then evaporated off in a vacuum concentrator centrifuge for 4 h, and the residue was reconstituted in 100 μL of methanol. Then, the supernatants were transferred to appropriate vials for determination of the concentration of LTB_4_ by a competition enzyme immunoassay, according to manufacturer’s instructions (EIA, Amershan Biosciences, Piscataway, NJ, USA). Quantification was accomplished using calibration curve containing LTB_4_ synthetic standards (Cayman Chemical, Ann Arbor, MI, USA).

### Macrophage cell culture (J774.1)

The J774.1 murine macrophage cell line was obtained from the American Type Culture Collection (ATCC, Rockville, MD, USA). Cells were cultured in Dulbecco′s Modified Eagle′s Medium (DMEM) supplemented with 10% fetal bovine serum (FBS) and 1% Penicilin/Streptomicin (Gibco, Grand Island, NY). After the formation of a monolayer, cells were harvested with plastic cell scrapers and centrifuged at 1,500 rpm for 10 min at 10°C. Next, supernatants were discarded and 10 mL of DMEM was added to each tube of cells. Cell viability and total cell numbers were determined by counting live and dead cells in a Neubauer chamber (BOECO Germany, Hamburg, Germany) after staining with Trypan blue (Gibco). Cells were plated in 96-well culture plates (Cell Wells – Corning Glass Workers) at a density of 1x 10^5^ cells / well and incubated overnight in DMEM in an incubator with a moist atmosphere of 5% CO_2_ at 37° C.

### Treatment with immunostimulatory and pharmacological agents

For osteoclastic cell differentiation, soluble mediator RANKL and monocyte colony stimulating factor (M-CSF / CSF-1) were added to the medium. Briefly, cells were cultured in the presence of M-CSF (30 ng / mL; R&D Systems, Minneapolis, USA), and with RANKL (10 ng / mL; R&D Systems). To confirm the osteoclastic phenotype, the cells were fixed and marked for TRAP (tartrate resistant acid phosphatase). The expression of genes that indicate an osteoclast phenotype was performed by RT-PCR in real time. Then, cells were plated at a density of 1 × 10^5^ cells per well in 96-well culture plates and stimulated with LTB_4_ encapsulated in microspheres at 0.01 µM and 0.1 µM, for 12, 24, 48 and 72 hours. For experimentation, DMEM medium without FBS was used and the plates were kept in an incubator at 37° C and 5% CO_2_.

### Cytotoxicity – Lactate dehydrogenase (LDH) assay

For cytotoxicity assessment, cells were plated in serum-free medium, at a concentration of 1 × 10^5^ cells per well, in 96-well plates and kept in an incubator at 37°C and 5% CO_2_ for 12 hours (*overnight*). After this period, cultures were stimulated with different concentrations of pharmacological and immunostimulating agents, for 24 hours. Next, 50 µL of the supernatant was collected and transferred to a new 96-well plate with a transparent, flat bottom and 50 µL of the CytoTox 96® Reagent was added to each sample. The plate was then covered with foil to protect against light and the samples incubated at 25° C for 30 minutes. After this period, 50 μL of the Stop Solution was added to each well. The absorbance was measured at 490 nm with a spectrophotometer (mQuanti, Bio-Tek Instruments, Inc., Winooski, VT, USA). As positive control, 10× Lysis Solution was added to the cells, 45 minutes prior to adding CytoTox 96® Reagent. LDH levels were expressed as percentages, according to the formula: cytotoxicity (%) = 100 × Experimental LDH Release absorbance / Maximum LDH Release absorbance (positive control). Groups were compared using the one-way ANOVA test followed by Dunnett’s post-test (α = 0.05).

### Determination of osteoclast formation by means of the activity of tartrate-resistant acid phosphatase (TRAP) enzyme

Cells were incubated in a solution containing 8 mg of naphthol AS-MX di-sodium phosphate salt (Sigma-Aldrich) in 500 µL of NN-dimethylformamide, followed by the addition of 50 mL of a 0.2 mol / L buffer solution sodium acetate (pH 5.0), containing 70 mg of Fast Red ITR (Sigma-Aldrich). Then the substrate sodium tartrate dihydrate (50 mmol / L) was added to the solution and incubated at 37° C for 2 hours. Subsequently, the wells were washed in distilled water and the cells stained with Harris’ hematoxylin. As a control, cells were incubated with medium without the substrate. The analysis of the formation of osteoclasts positive for the TRAP enzyme was determined based on the presence or absence of labeling, in two independent experiments. First, the plates were photographed under bright field microscopy, in 10 × magnification. For quantification of multinucleated TRAP + cells by field of view, the Software Image J (National Institutes of Health, Bethesda, MD, USA) and the image deconvolution plugin (Color Deconvolution) were used. Fast Red vector was applied and then the selected red channel and threshold were manually adjusted. To count the number of cells, the “Analyze Particles” tool was used, after calibration so that the count included only structures with a minimum pixel size of 500, which is equivalent to the size of a multinucleated cell with at least 3 nuclei. The groups were compared using the one-way ANOVA test followed by the Tukey post-test (α = 0.05).

### RNA extraction, reverse transcription, and polymerase chain reaction in real time (qRT-PCR)

mRNA levels were measured by quantitative reverse transcriptase-polymerase chain reactions (qRT-PCR) after cell stimulation. To this end, total RNA was extracted using the RNeasy® Mini kit (Qiagen Inc., Valencia, USA) and quantified using NanoDrop 2000 spectrophotometer (Thermo Fisher Scientific Inc., Wilmington, USA). A total of 1 µg of RNA were used for cDNA synthesis with the High Capacity cDNA Reverse Transcription kit (Applied Biosystems, Foster City, USA) in a thermal cycler (Veriti® Thermal Cycler, Applied Biosystems, USA). qRT-PCR reactions were performed in duplicate using the TaqMan® system in a StepOne Plus® real-time PCR system (StepOne Plus® Real-Time PCR System, Applied Biosystems) and the following cycle program: 95 °C for 20 s, 40 cycles at 95 °C for 1 s, and 60 °C for 20 s. Primer-probe pairs included *Alox5* (Mm01182747), *Alox5ap* (Mm00802100), *Acp5* (Mm00475698), *Mmp9* (Mm00442991), *Ctsk* (Mm00484039) and *Calcr* (Mm01197736)(TaqMan**®** Gene Expression Assay, Applied Biosystems). All protocols were performed according to the manufacturers’ instructions. Glyceraldehyde-3-phosphate dehydrogenase (*Gapdh*) and beta-actin (*Actb*) were used as reference genes for normalization purposes. The results were analyzed based on cycle threshold (Ct) values. Relative expression was calculated by the ΔΔCt method. Groups were compared using one-way ANOVA test followed by the Tukey post-test or the Dunnett post-test (α = 0.05).

## Results

### 5-LO pathway is involved in the osteoclastic differentiation of cells of the hematopoietic lineage

During osteoclast differentiation induced by M-CSF and RANKL, 5-lipoxygenase gene expression (*Alox5*) was induced 48 hours after the addition of the stimulus (p <0.05), as well as the expression of the activating protein of 5-lipoxygenase (*Alox5ap*), induced 48 and 72 hours later (p <0.05) (Figure 1). These results indicate that the 5-LO pathway and the metabolites produced during the arachidonic acid metabolism may be involved in the osteoclast differentiation process.

**Figure 1.**
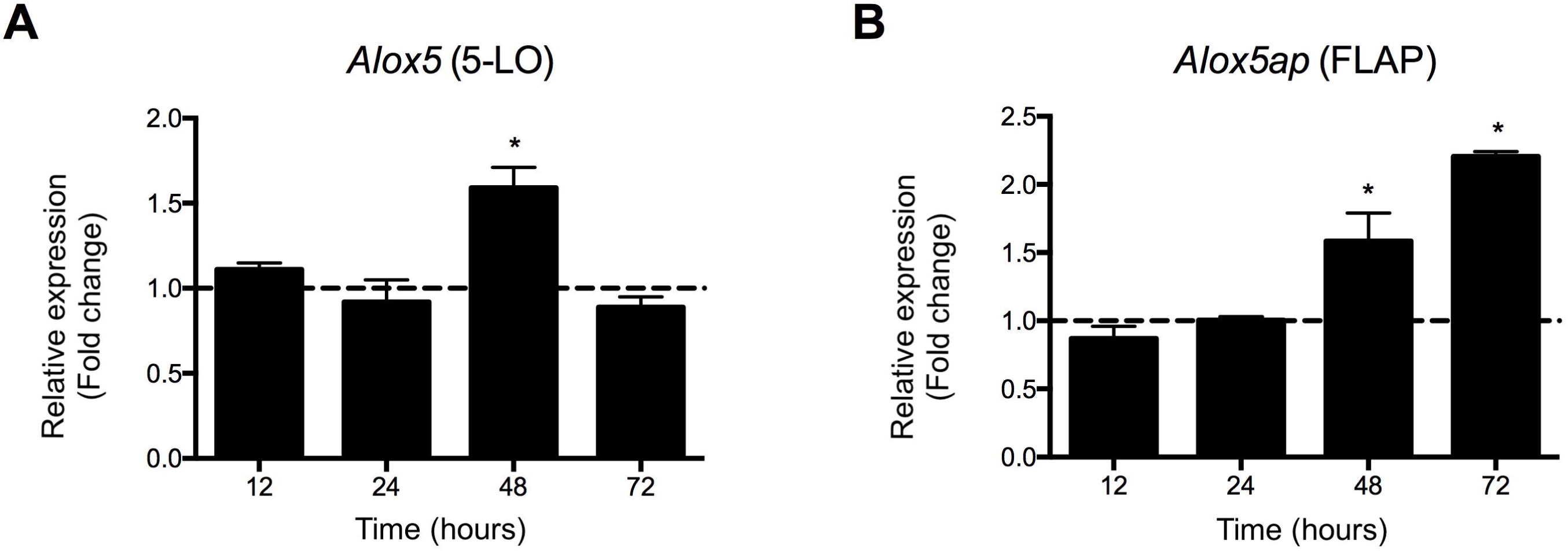
Relative expression of *Alox5* (A) and *Alox5ap* (B), genes which encode 5-LO enzyme and the 5-LO activating protein (FLAP), respectively, 12, 24, 48 and 72 hours after stimulation of murine macrophages J774A.1 with M-CSF (30 ng / ml) and RANKL (10 ng / ml). * p <0.05 compared to cells maintained in culture medium without serum (dashed line).

### Characterization of microspheres prepared for sustained delivery of LTB_*4*_

No bacterial contamination was found in any batch of prepared microspheres, after culture in BHI-agar medium, for a period of 24 hours in an incubator. Likewise, no endotoxin was detected in the samples of each prepared batch. Therefore, empty microspheres or containing encapsulated LTB_4_ were used in subsequent experiments.

After lyophilization of the microspheres obtained by the solvent evaporation method, the characterization was carried out by dispersing their diameters, after reconstitution in distilled and deionized water. The average diameter of the empty microspheres was 4.1 ±2.7 μm (Figure 2A) and the microspheres containing LTB_4_ (MS-LTB_4_) was 5.1 ±4.5 μm (Figure 2B). Therefore, the microspheres did not have their diameters changed by the LTB4 encapsulation process.

**Figure 2.**
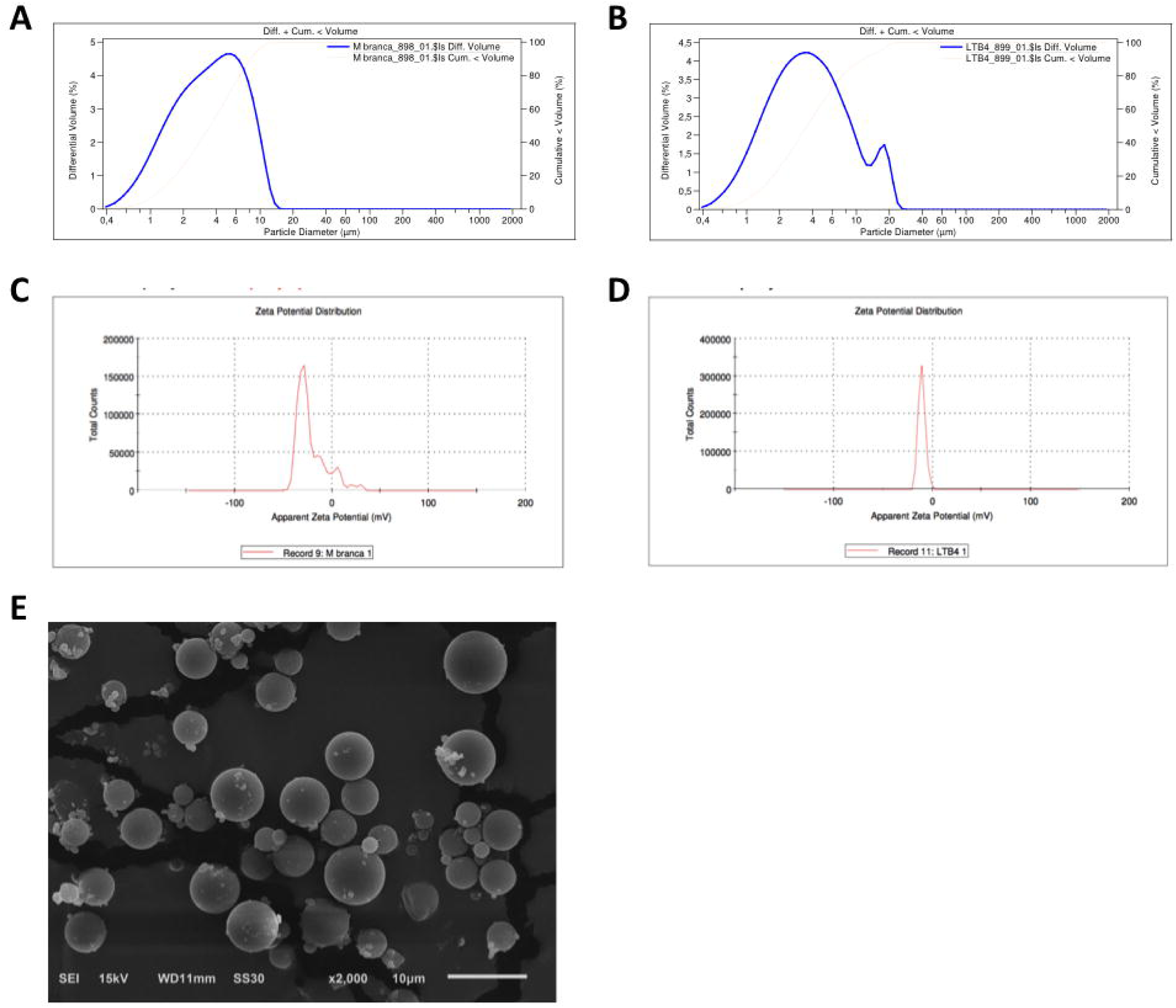
Characterization of empty microspheres (A, C, E) or containing LTB_4_ (B, D, F), according to their size, zeta potential and morphology. Diameter distribution in μm (A, B), distribution of the zeta potential in mV (C, D) and topography in scanning electron microscopy (E, F).

Regarding the electrical stability of the microspheres, there were no relevant differences between the means of the zeta potential of the empty microspheres (−21.7mV ± 5.5 mV (Figure 2C) and of the microspheres containing LTB_4_ (MS-LTB_4_) (−10.7mV ± 3.6 mV, Figure 2D) Therefore, LTB_4_ encapsulation did not result in changes in the electrical charge on the PLGA surface.

Shape and topography of the microspheres were determined using scanning electron microscopy. Overall, the microspheres empty or containing LTB_4_ (MS-LTB_4_) presented spherical shape, uniform surfaces and without pores (Figure 2E and 2F).

### Exogenous addition of LTB4 embedded in microspheres inhibits osteoclastogenesis

Because we observed a positive modulation of the 5-LO pathway during osteoclastogenesis, we investigated whether the exogenous addition of LTB_4_ could modify the course of cell differentiation. For this, LTB_4_ was used in two different concentrations (0.01 μM and 0.1 μM), incorporated in microspheres of lactic acid-co-glycolic acid (PLGA). This strategy was used because the osteoclast differentiation experiment is long (12 to 72 hours) and the half-life of the lipid mediators is short. Thus, we aimed to keep LTB_4_ available for the entire maintenance period of the cell culture. The encapsulation efficiency of LTB_4_ in PLGA microspheres was 42%.

Microspheres were not cytotoxic to macrophages, regardless of whether they contained LTB_4_ or not, compared to DMSO (positive control; p <0.05). The stimulation with M-CSF + RANKL was not cytotoxic compared to the DMEM culture medium (p> 0.05) (Figure 3).

**Figure 3.**
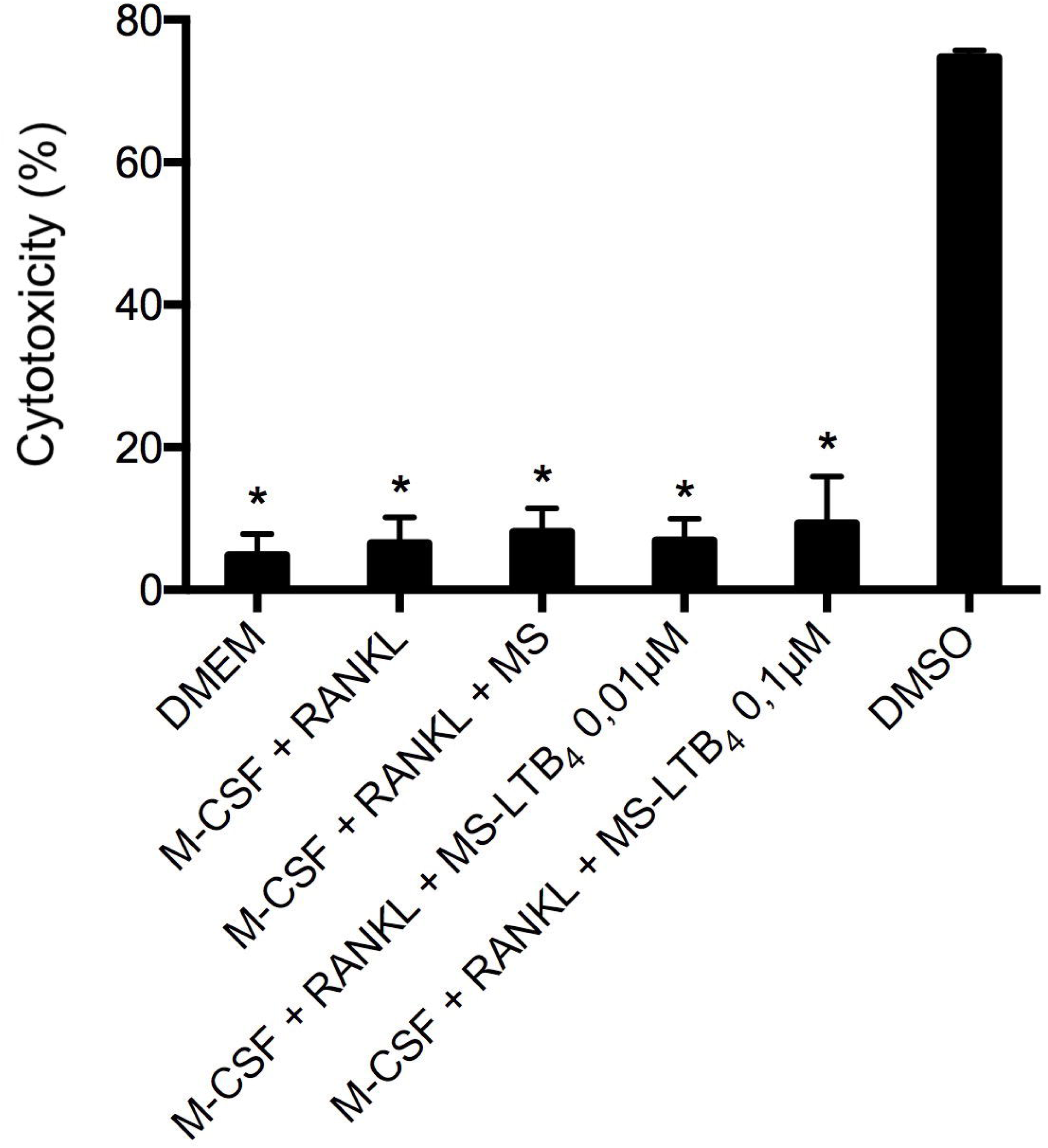
Cytotoxic effect of 0.01 μM and 0.1 μM of LTB_4_ encapsulated in microspheres and empty microspheres, as well as M-CSF + RANKL stimuli, in culture of murine macrophages J774A.1, measured by through the LDH test, 24 hours after exposure. * p <0.05 compared to the positive control (DMSO).

After 72 hours, treatment with M-CSF + RANKL led to the formation of cell colonies and differentiation of TRAP positive osteoclasts, differently from what was observed in the group maintained with DMEM medium without FBS. Interestingly, treatment with M-CSF + RANKL + LTB4 microspheres led to less osteoclast formation on the culture plate, regardless of the concentration used (p <0.05). The treatment with M-CSF + RANKL + empty microspheres showed a result similar to M-CSF + RANKL (p> 0.05). These results show that LTB_4_, a metabolite of the 5-LO pathway produced by the action of the leukotriene A4 hydrolase enzyme on LTA_4_, reduces the osteoclastogenic potential induced by M-CSF + RANKL (Figure 4).

**Figure 4.**
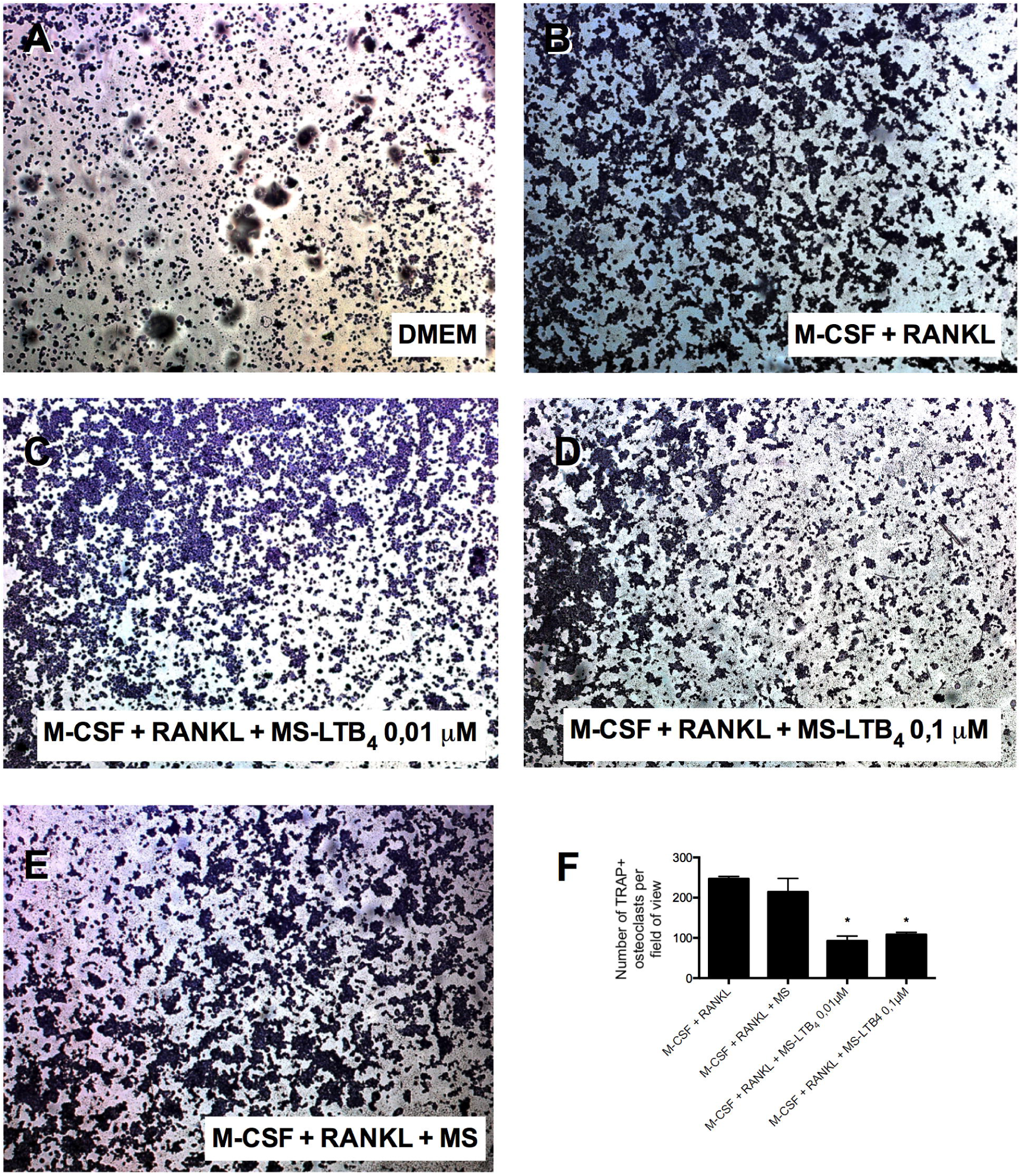
Representative images of J774A.1 murine macrophages and differentiated osteoclasts (TRAP +), cultured in DMEM (control) medium (A), with M-CSF (30 ng / mL) + RANKL (10 ng / mL) (B), with M-CSF (30 ng / mL) + RANKL (10 ng / mL) containing microspheres (MS) of LTB_4_ in concentrations of 0.01 μM (C) and 0.1 μM (D), and with M-CSF (30 ng / mL) + RANKL (10 ng / mL) containing empty microspheres (MS) (E), 72 hours after adding the stimuli. Graph shows the quantification of the number of osteoclasts, by field of view, in the 5 × (F) magnification.

### LTB4 induces Mmp9 expression but inhibits Calcr and Ctsk, without changing Acp5

In order to investigate the effect of exogenous addition of LTB4 on the synthesis of molecules involved in osteoclast differentiation, the gene expression of a panel of markers of clast formation and activity was evaluated.

*Acp5* gene, which encodes the TRAP enzyme, was not modulated by the addition of M-CSF + RANKL with or without LTB_4_, 48 or 72 hours after the stimulus (p> 0.05). The *Mmp9* gene, which encodes matrix-9 metalloproteinase enzyme, was stimulated by the addition of M-CSF + RANKL to the culture medium, both 48 and 72 hours after stimulation (p <0.05). Interestingly, after 48 hours LTB_4_ had no effect on the synthesis of *Mmp9* (p> 0.05), but after 72 hours there was a greater production of *Mmp9* after the addition of 0.1 μM or 0.01 μM LTB_4_ (p < 0.05). Of these, the effect was more pronounced with 0.01 μM LTB_4_(p <0.05) (Figure 5).

**Figure 5.**
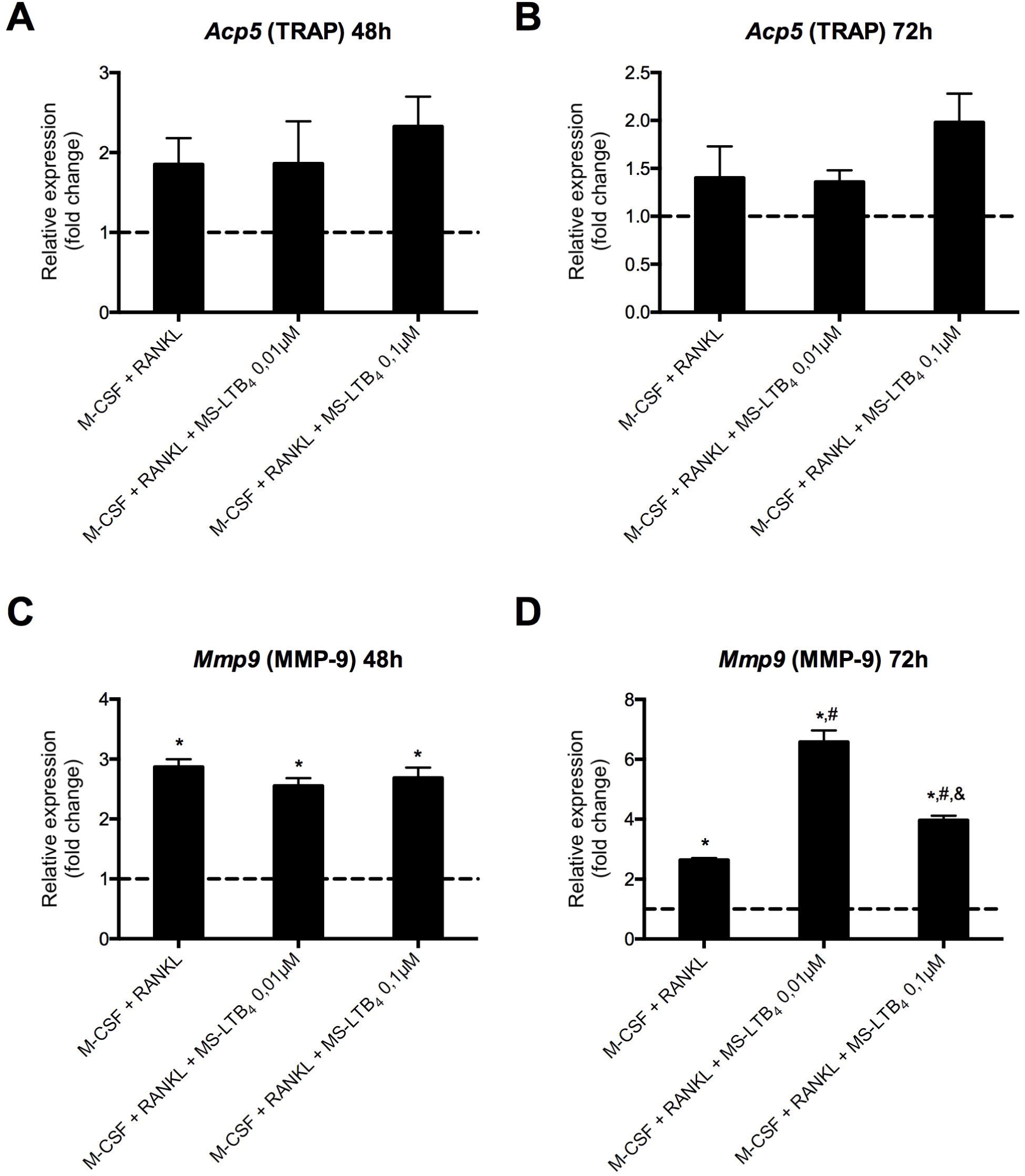
Relative expression of the *Acp5* (A, B) and *Mmp9* (C, D) genes that encode the TRAP and matrix-9 metalloproteinase enzymes respectively, 48 and 72 hours after stimulation of murine macrophages J774A.1 with M-CSF (30 ng / mL) + RANKL (10 ng / mL) containing or not LTB_4_ microspheres (MS) at concentrations of 0.01 μM and 0.1 μM. * p <0.05 compared to expression in cells maintained in DMEM medium without fetal bovine serum (dashed line); # p <0.05 compared to adding M-CSF + RANKL only; & p <0.05 comparing the different LTB_4_ concentrations.

*Ctsk* gene, which encodes the enzyme cathepsin K, was not modulated by adding M-CSF + RANKL to the culture medium (p> 0.05), but the expression was decreased after adding 0.01 μM LTB_4_ (p < 0.05), both at 48 and 72 hours. *Calcr* gene, which encodes the calcitonin receptor, showed a different pattern depending on the period evaluated. After 48 hours, M-CSF + RANKL induced its expression (p <0.05) and the addition of LTB_4_ at 0.01 μM or 0.1 μM inhibited expression until it returned to baseline (p <0.05). Differently, after 72 hours, the addition of LTB_4_ at 0.01 μM or 0.1 μM did not influence the expression induced by M-CSF (p> 0.05) (Figure 6).

**Figure 6.**
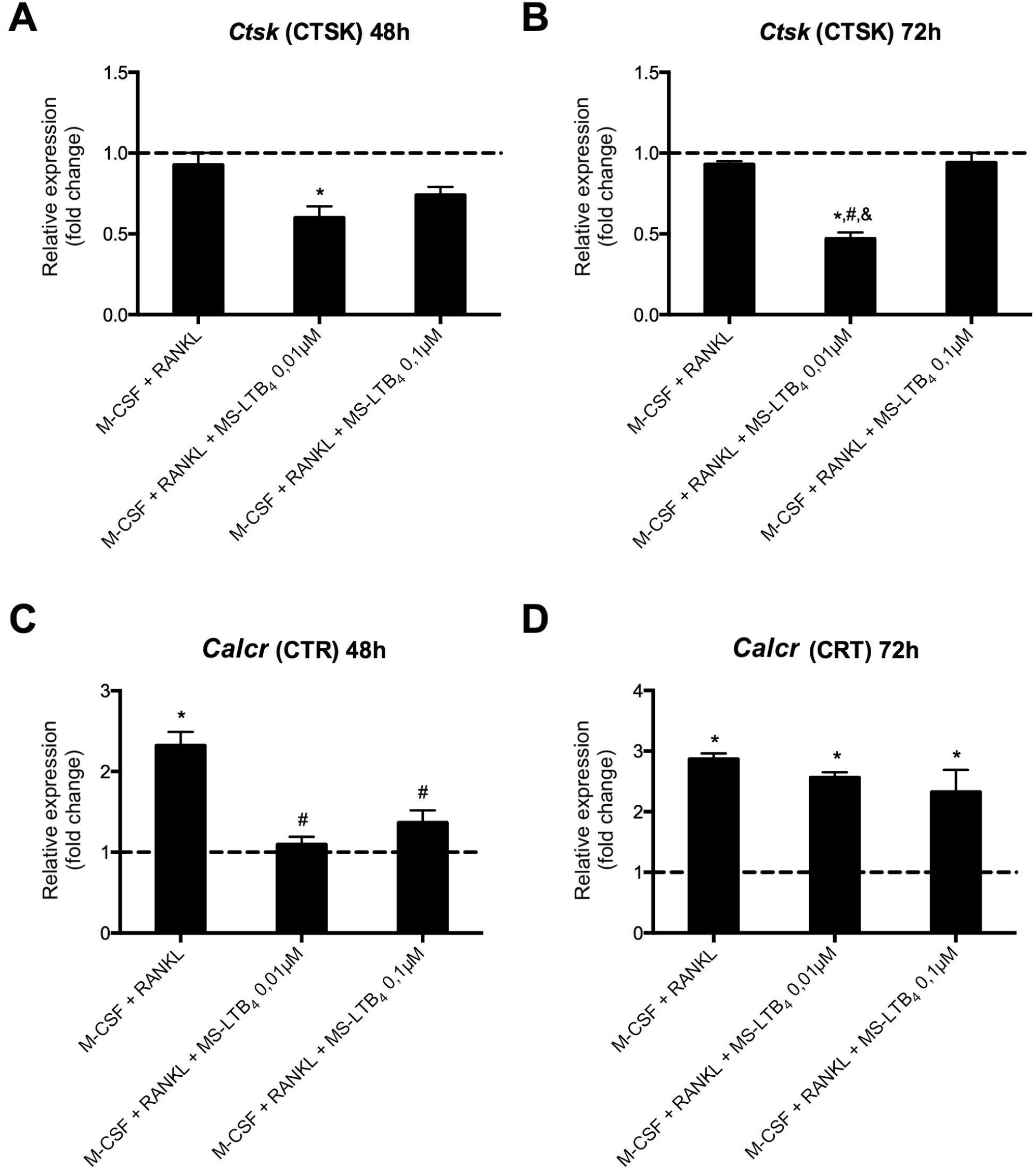
Relative expression of *Ctsk* (A, B) and *Calcr* (C, D) genes encoding cathepsin K and calcitonin receptor respectively, 48 and 72 hours after stimulation of murine macrophages J774A.1 with M-CSF (30 ng / mL) + RANKL (10 ng / mL) containing or not LTB_4_ microspheres (MS) in concentrations of 0.01 μM and 0.1 μM. * p <0.05 compared to expression in cells maintained in DMEM medium without fetal bovine serum (dashed line); # p <0.05 compared to adding M-CSF + RANKL only; & p <0.05 comparing the different LTB_4_ concentrations.

## Discussion

Modulation of the 5-LO proinflammatory pathway was observed *in vitro* during hematopoietic cell differentiation into osteoclasts and the addition of LTB_4_-containing microspheres inhibited the osteoclastogenic potential of the culture induced by the soluble mediator RANKL. These results corroborate with our *in vivo* observations that this pathway plays a protective role during *in vivo* osteoclastogenesis (Paula-Silva et al., 2016; Paula-Silva et al., 2020).

Inflammatory process in the face of aggressive microbial stimuli is a response from the host that aims to fight infection and restore its physiological functioning (Gilroy et al., 2010). Cytokines and various signaling molecules are synthesized and secreted by resident cells, prior to the recruitment and activation of cells of the immune system, generating a molecular pattern of immune response (Cooper et al., 2011). In this process, lipid mediators derived from arachidonic acid are important molecules in the regulation of homeostatic and inflammatory processes (Dennis and Norris, 2015).

In bone tissue, there is a constant interrelation between the inflammatory response and the resorption of mineralized matrices. Thus, considering that we observed a positive modulation of the 5-LO pro-inflammatory pathway during osteoclastogenesis and that a previous study showed that LTC_4_ is important for osteoclastic differentiation of bone marrow-derived macrophages (Lee et al., 2012), we investigated whether LTB_4_ added exogenously could modify the course of cell differentiation and clarify a possible mechanism involved in the periapical response observed in animals deficient in the 5-LO enzyme.

Microspheres were used as the delivery system for LTB_4_ in this study. This encapsulation method allows LTB_4_ to maintain its biological activity preserved, in addition to being an important tool for cellular stimulation, in order to protect the encapsulated mediator from degradation (Nicolete et al., 2008). After evaluation for a period of 24 hours, it is known that microspheres of LTB4 had a peak release of 45% of the mediator in a period of 5 hours, and can be continuously released for a long period of time (Nicolete et al., 2007). The microspheres were prepared using a polymer, the co-glycolic poly-lactic acid (PLGA) as a scaffold. PLGA is widely used for preparation of microspheres because it is a biodegradable and biocompatible polymer (Fredenberg et al., 2011; Pereira et al., 2015; Reis et al., 2017; Lorencetti-Silva et al., 2019). This material allows the release of encapsulated molecules in a controlled manner for long periods and from a single administration, which keeps the concentrations of the encapsulated substance constant (Ford Versypt et al., 2012).

LTB_4_ has been efficiently encapsulated in PLGA previously (Nicolete et al., 2007; Nicolete et al., 2008), without prejudice to its biological activity. This polymer has high encapsulation efficiency, stability and its degradation products (lactic and glycolic acid) are also biocompatible (Bitencourt et al., 2015; Pereira et al., 2015). Specifically with respect to LTB_4_, this delivery system proves to be a relevant alternative to conventional drug delivery systems, since in addition to releasing the encapsulated molecules in a controlled manner over longer periods through a single administration, it maintains the stability of mediators extremely labile (Ford Versypt et al., 2013; Pereira et al., 2015). Lipid mediators can be easily degraded, via oxidation and hydrolysis reactions, and consequently lose their biological properties (Murphy and Gijón, 2007). Therefore, encapsulation is an efficient way to maintain the constant concentration of lipid mediators, in order to prevent its degradation.

In a previous study, LTB_4_ induced RANKL expression, which in turn led to osteoclast differentiation (Chen et al., 2010). In the absence of metabolites from the 5-LO pathway, there was a reduction in recruitment and differentiation of osteoclastic cells, preventing bone resorption induced by mechanical load (Moura et al., 2014). However, a comparison of these studies regarding the osteoclastogenic potential of LTB_4_ should consider some factors. The first is that this modulation can be modified by the presence or absence of concomitant infection, as well as by the concentration of the mediator used. The second, and perhaps more relevant, is the delivery system used. Up to date, the role of LTB_4_ in osteoclastogenesis was evaluated using the mediator in the soluble form (Jiang et al., 2005; Chen et al., 2010; Lee et al., 2012; Dixit et al., 2014), which acts mainly on the BLT1 and BLT2 surface receptors. We used the encapsulation in PLGA microspheres as a strategy. Encapsulation favors the engulfment of microspheres by phagocytes such as macrophages J774A.1 (Bitencourt et al., 2015), which allows this mediator to be released inside the cell and signal via nuclear receptors of the PPAR family. Polymeric microspheres containing LTB_4_ are more phagocytized by murine peritoneal macrophages than empty microspheres (Nicolete et al., 2007; Nicolete et al., 2008) and are capable of inducing an increase in the production of nitric oxide in human endothelial cells and in peritoneal macrophages murine, and increase the expression of the nuclear receptor PPAR-α (Nicolete et al., 2008). This mechanism has not been demonstrated in our work, but we believe it can be important and should be explored in future studies in order to understand the dual role of the lipid mediator, dependent on the family of receptors that are activated in the signaling cascade.

Stimulation of human mononuclear cells with M-CSF associated with LTB_4_ increased the expression of osteoclastogenesis marker genes such as *CTSK, MMP9* and *ACP5* (Dixit et al., 2014), differently from what we observed with an immortalized culture of macrophages obtained from mice, in which LTB4 induced the expression of *Mmp9* but inhibited *Calcr* and *Ctsk*, without changing *Acp5*. These divergent results could be achieved due to the incorporation of LTB_4_ in PLGA microspheres and shed light to a novel protective role of the lipid mediator as inhibitor of osteoclast differentiation.

## Conclusion

Leukotriene B4 encapsulated into microspheres inhibited differentiation of macrophages into an osteoclastic phenotype and activation of the cells under RANKL stimulus.

## Acknowledgements

This study was supported by Grants from São Paulo Research Foundation (FAPESP) 2010/17611-4 to FWGPS and 2014/14015-2 to JPQT. FLS received a CAPES Fellowship (Financial Code: 000).

